# Reorienting Spatial Attention within Visual Working Memory

**DOI:** 10.1101/854703

**Authors:** Sizhu Han, Yixuan Ku

**Author notes:** correspondence: Yixuan Ku.

## Abstract

Attention and working memory (WM) are intertwined core cognitive processes. Through four experiments with 133 participants, we dissociated the impact of two types of covert spatial attention, endogenous *vs.* exogenous, on visual WM. Behavioral results consistently indicated that exogenous attentional cues were more advantageous than endogenous ones in enhancing the precision of visual WM under load-2, while they equalized under load-4. In addition, physiological and neural data explained the mechanisms. Converging evidence from eye-tracking, electroencephalography, and magnetoencephalography suggested that fast attentional processing induced by exogenous cues lead to early top-down information from the dorsal lateral prefrontal cortex (DLPFC) to sensory cortices. The differential frontal activities were further correlated with the behavioral distinctions between exogenous and endogenous cues, and transcranial magnetic stimulation over DLPFC at the same time period abolished the exogenous advantage. Taken together, traditionally considered bottom-up attentional processing induced by exogenous cues rapidly engages top-down signals from the frontal cortex, which leads to stronger behavioral benefits compared with the benefits produced by endogenous cues under the low load condition.

## INTRODUCTION

Working memory (WM) is a critical ability to store and manipulate sensory information when it is no longer accessible in the environment (Baddeley, 1986). Visual working memory (VWM) capacity is severely limited (Cowan, 2001; Miller, 1956), but lies in the center of cognition and is linked to the general intelligence (Engle, Kane, & W Tuholski, 1999). VWM representations, however, are flexible (Van den Berg, Awh, & Ma, 2014) and could be influenced by retro-cues (Griffin & Nobre, 2003; Souza & Oberauer, 2016), indicating interactions between attention and VWM.

As complex as memory, attention also has many different sub-types from distinct standards of classification (Cowan, 2017; Carrasco, 2011; Fan, McCandliss, Fossella, Flombaum, & Posner, 2005; Posner, 1980). Since William James, attention has been divided into two types: one that is reflexive and involuntary, and another that is active and voluntary (James, Burkhardt, Bowers, & Skrupskelis, 1890; Johnston & Dark, 1986). These two kinds of attention correspond to bottom-up and top-down attention from neural views (Buschman & Miller, 2007; Hahn, Ross, & Stein, 2006; Theeuwes, 2010). Covert attention, as opposed to overt attention, is the attentional deployment towards peripheral locations while the eyes gaze at the center. It can be directed by exogenous or endogenous cues, corresponding to the above reflexive/involuntary/bottom-up and active/voluntary/top-down ways respectively.

In the perception domain when the attentional cues are given prior to the sensory stimuli, a plethora of studies indicated that the exogenous attention processes rise (∼100ms) and decay quickly (Liu, Stevens, & Carrasco, 2007; Remington, Johnston, & Yantis, 1992), while the endogenous attention processes deploy slower (∼300ms) but remains sustained (Busse, Katzner, & Treue, 2008). However, the two attentional effects over perception are normally equivalent given enough time for attention to deploy (See review in Carrasco, 2011), although there exist part of evidence claiming their behavioral difference when sensory signal-to-noise ratio is high (Lu & Dosher, 1998, 2000; Lu, Lesmes, & Dosher, 2002). Despite of their similar behavioral benefit, the two types of cues have already been suggested to induce neural processes that involve overlapped but distinctive neural networks (Buschman & Miller, 2007) (Chica, Bartolomeo, & Lupiáñez, 2013; Carrasco, 2011; Corbetta & Shulman, 2002), supporting the dissociated neural view of bottom-up vs. top-down.

Meanwhile, in the memory domain when the attentional cues are given during the delay period after the sensory stimuli disappear, many studies have suggested that the two attentional effects over memory representations can enhance behavioral performance (e.g., Astle, Summerfield, Griffin, & Nobre, 2012; Brady & Hampton, 2018; Gözenman, Tanoue, Metoyer, & Berryhill, 2014; Gunseli, van Moorselaar, Meeter, & Olivers, 2015; Lepsien & Nobre, 2006; Murray, Nobre, Clark, Cravo, & Stokes, 2013; Williams, Hong, Kang, Carlisle, & Woodman, 2013; Griffin & Nobre, 2003; Matsukura, Cosman, Roper, Vatterott, & Vecera, 2014; Pertzov et al., 2013; Tanoue & Berryhill, 2012). However, none of them have found significant difference between them. There might be several reasons for such null effects. First, insensitive binary report with a change detection task (Shimi, Nobre, Astle, & Scerif, 2014) may not catch their subtle difference. Second, relatively high task difficulty (load 4) with a free-recall task (Pertzov et al., 2013) is similar to the condition of low signal-to-noise ratio in the perception domain. Increasing the memory signal-to-noise ratio might reveal the distinctions in the memory domain, given several findings of the differences in the perceptual domain when the sensory-to-noise ratio is high (Lu & Dosher, 1998, 2000; Lu, Lesmes, & Dosher, 2002). Third, the effects of endogenous and exogenous cues in the memory domain are truly similar after excluding the above two reasons. Nevertheless, even if the endogenous and exogenous cues produce similar behavioral results in the memory domain, they may induce different neural processes and involve different neural networks, seeing that neural networks induced by them in the perceptual domain are distinctive (Buschman & Miller, 2000; Chica, Bartolomeo, & Lupiáñez, 2013; Carrasco, 2011; Corbetta & Shulman, 2002) and previous studies indicate similarities between the neural circuits implemented in orienting attention towards representations in perception and memory (Awh & Jonides, 2001; Heuer & Schubö, 2016; Lepsien & Nobre, 2006; Myers, Walther, Wallis, Stokes, & Nobre, 2015).

Altogether, it urges experimental designs with more sensitive task, as well as varied levels of task difficulty, to reveal the potential difference between the endogenous and exogenous attentional effects in the memory domain, in such a way we could unify the attentional effects in both perception and memory. Moreover, neuroimaging methods with high temporal resolutions are needed to catch the potential difference in neural processes between them. Therefore, we conducted 4 experiments, combined with physiological and neural measurements, including eye-tracker, electroencephalography (EEG), magnetoencephalography (MEG), and transcranial magnetic stimulation (TMS), to address the following two questions: 1) whether there is a behavioral or neural difference between the endogenous and exogenous attentional effects in the memory domain? 2) If there exists a difference, when and how the endogenous and exogenous attentional effects differentiate?

## RESULTS

### Experiment 1

In the first experiment, a total of 64 participants were requested to memorize two or four oriented Gabor patches (presented for 500 ms) that differed by at least 10 degrees. After a delay of 2 s, one of the Gabor patches was probed. The participants reproduced its orientation using a computer mouse as precisely as possible. In two thirds of all the experimental trials, a retro-cue was presented in the middle of the 2 s delay. Half of the retro-cue trials used an endogenous cue (i.e. an arrow at the center of the screen) and the other half used an exogenous cue (i.e. a dashed circle outside of one of the Gabor patches). All cues were 100% valid, indicating the upcoming probed location. All trials were randomly presented. To control eye movements and blinks, gaze positions were on-line monitored during task performance (see Methods for more details).

As the best fitting model of memory recall task were still in debate (Adam, Vogel, & Awh, 2017; Ma, Husain, & Bays, 2014), we mainly focused on reporting the results from the raw circular standard deviation of the recall error (i.e. Raw SD). Six participants in experiment 1 were excluded from further analysis due to either poor performance (lower end in the group 99% confidence interval, the same below) in at least one condition (4 participants) or data missing (2 participants). Raw SD from the rest of 58 subjects was shown in Fig. 2a. Baseline-corrected pupillary diameters were calculated as an index to reflect physiological states (e.g. larger pupil size is associated with a higher level of arousal (Alnæs et al., 2014; Granholm & Steinhauer, 2004; Nassar et al., 2012)) over time. The pupillary diameters are illustrated in Fig. 3a-b. We performed two-way repeated measures ANOVAs with the within-subject factors of LOAD (2 and 4) and ATTENTION (endogenous, exogenous, and no-cue) for behavioral results. Post-hoc two-tailed t-tests were performed to examine the effect of ATTENTION under each load, and multiple comparisons were corrected using the false discovery rate (FDR) correction (the same for the following experiments).

**Figure 1.**
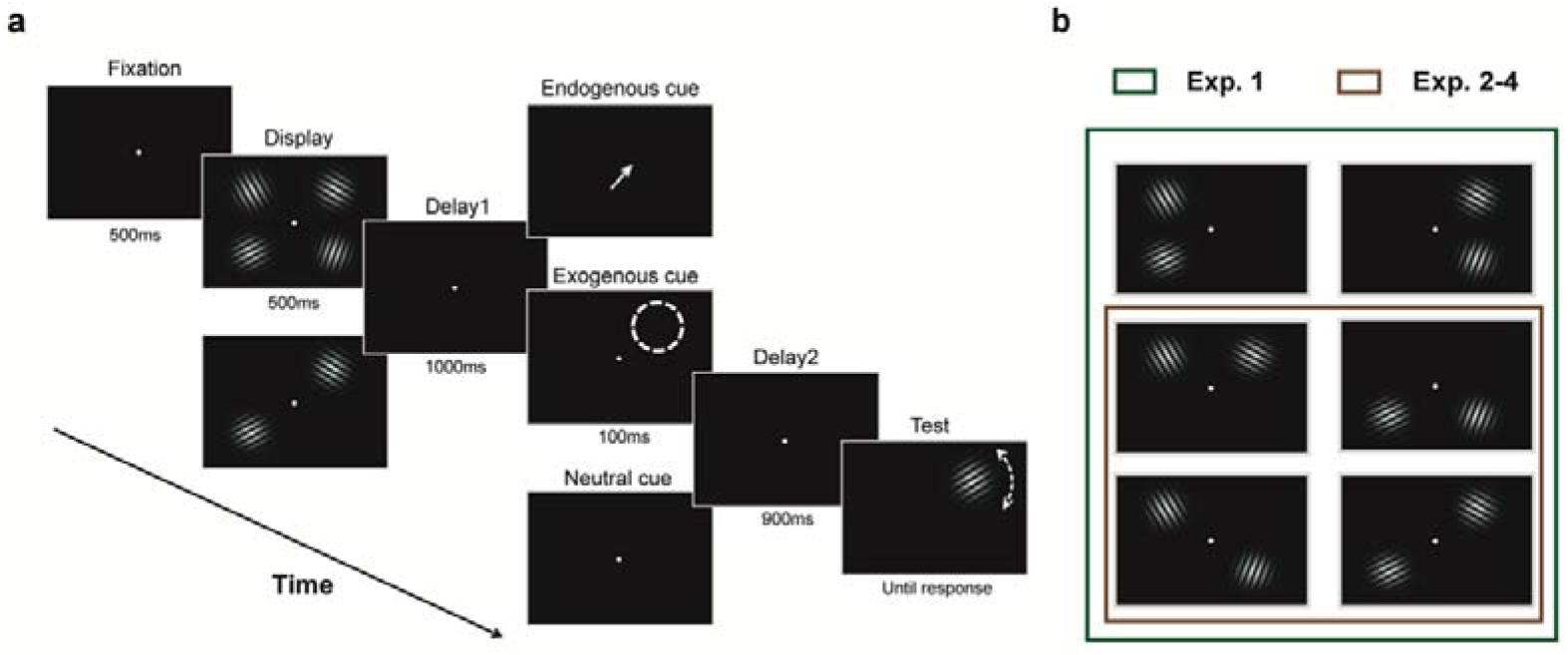
Trial sequence. **a**, Example of two types of valid retro-cue trials (endogenous and exogenous cue) and of a no cue trial. **b,** Configurations of the display that was set to load-2 in four experiments.

**Figure 2.**
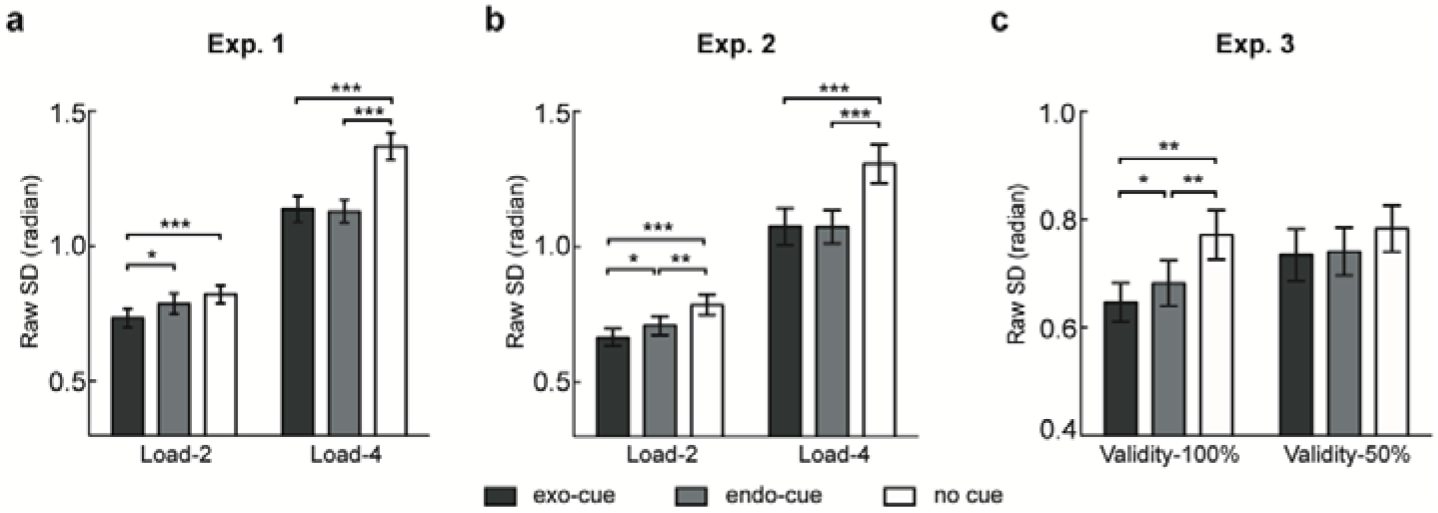
Behavioral results. **a**, Behavioral results from experiment 1 (*n*=58). **b,** Behavioral results from experiment 2 (*n*=19). **c,** Behavioral results from experiment 3 (*n*=24). * *p*<0.05, ** *p*<0.01, *** *p*<0.001, all *p*-values are FDR corrected.

**Figure 3.**
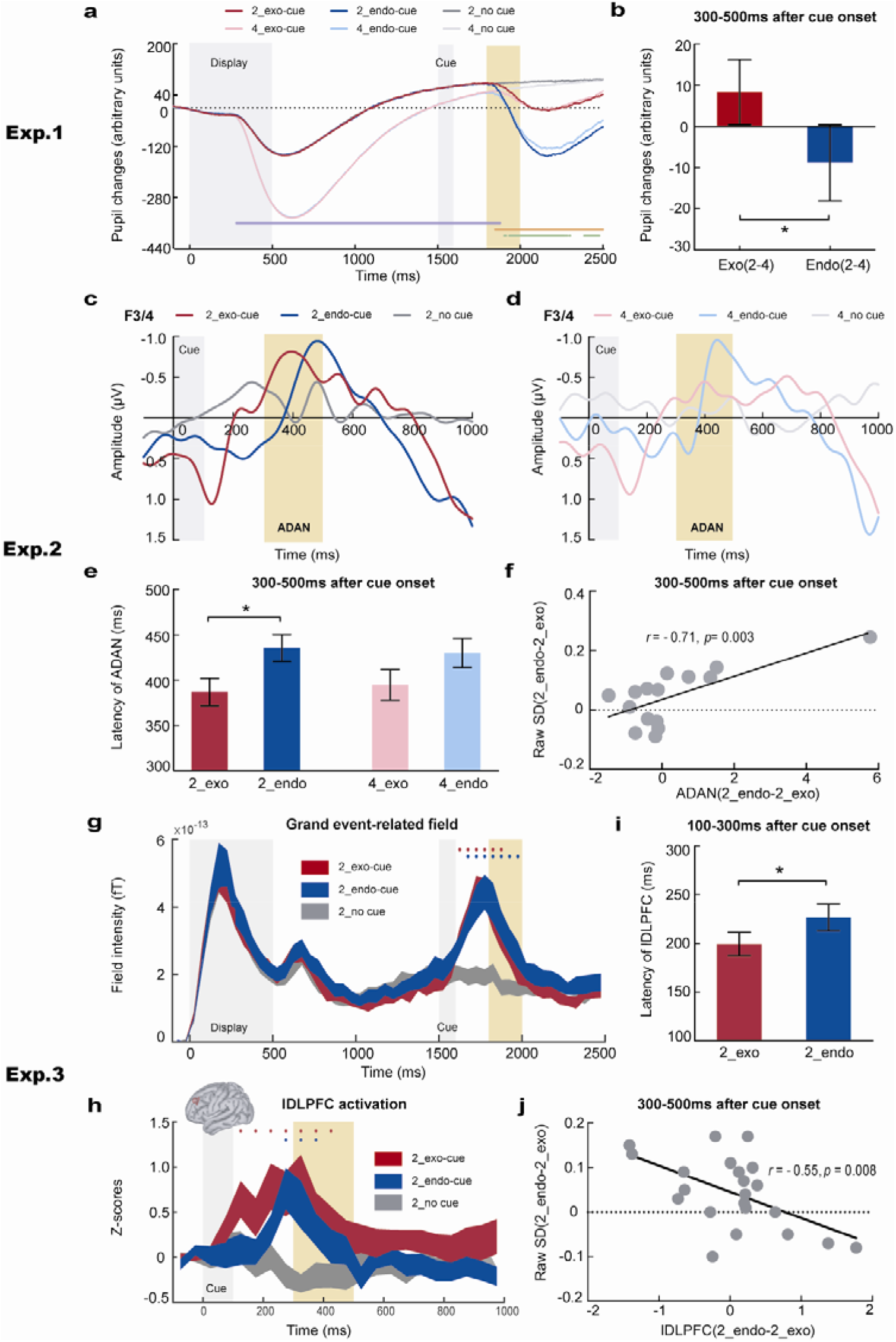
Psychophysiological results. **a**, Pupil changes as a function of time referred to the memory display. Purple dots indicate a main effect of LOAD, orange dots indicate a main effect of ATTENTION, and green dots indicate a 2-by-2 interaction. **b,** Load-by-cue interactions within the time window of 300-500 ms after the cue onset. **c-d,** The contra-minus ipsilateral curves at frontal F3/4 electrodes time-locked to the retro-cue onset for the load-2 (c) and load-4 condition (d). The yellow area indicates the time period of the ADAN component (300-500 ms after the cue onset). **e,** The latency of the ADAN component for retro-cue conditions. **f,** Correlations between ADAN and behavioral results under the load-2 condition (two-tailed *t*-tests, *p*<0.05). **g-h,** Grand event-related field time-locked to memory display (g) and the left DLPFC activation time-locked to the cue onset (h) in blocks with 100% validity. Red (blue) dots indicate higher brain responses to exogenous (endogenous) retro-cues than to no cue condition (one-tailed *t*-tests, *p*<0.05). Yellow areas indicate a time period of 300-500 ms after the cue onset. **i,** The latency of lDLPFC activation within the range of 100-300 ms after cue onset for retro-cue conditions. **j,** Correlations in blocks with 100% validity of activation differences between endogenous and exogenous cues at lDLPFC and their behavioral differences within a post-cue period of 300-500ms. * *p*<0.05.

2-by-3 ANOVAs on the Raw SD revealed that there were significant main effect of LOAD (*F*(1, 57) = 369.47, *p* =1.4e-26, η_p_^2^ = 0.87), significant main effect of ATTENTION (*F*(2, 114) = 46.21, *p* =2.0e-15, η_p_^2^ = 0.45), and a significant interaction between LOAD and ATTENTION (*F*(2, 114) = 26.40, *p* =3.8e-10, η_p_^2^ = 0.32).

Figure 2a displays the interaction. Under the load-4 condition, both endogenous and exogenous retro-cues facilitated VWM performance equivalently, compared to the no-cue condition (4_endo vs. 4_exo: *t(57)*=-0.38, *p_adj_*=0.71, Cohen’s *d*=0.05; 4_endo vs. 4_neu: *t(57)*=-8.20, p*_adj_*=4.8e-11, Cohen’s *d*=1.08; 4_exo vs. 4_neu: *t(57)*=-8.59, *p_adj_*=2.2e-11, Cohen’s *d*=1.13). Importantly, under the load-2 condition, exogenous cues were more effective than endogenous cues (2_endo vs. 2_exo: *t(57)*=2.64, *p_adj_*=0.017, Cohen’s *d*=0.35; 2_endo vs. 2_neu: *t(57)*=-1.70, *p_adj_*=0.094, Cohen’s *d*=0.22; 2_exo vs. 2_neu: *t(57)*=-5.82, *p_adj_*=8.5e-7, Cohen’s *d*=0.76).

To focus on the attentional processes induced by the retro-cues, 2-by-2 ANOVAs for the pupil diameters were performed at each time point. There existed significant main effects of LOAD starting from 284 ms after the sample stimuli until 376 ms after the cues (*p*s<0.05), significant main effects of ATTENTION (endogenous, exogenous) starting from 350 ms after the cues (*p*s<0.05), and significant interactions starting from 406 ms after the cues (*p*s<0.05). To further demonstrate this interaction, the sensory evoked activity could be subtracted between the two VWM loads within one attentional type. The difference between endogenous and exogenous attention was shown in Fig. 3b. Within the time window of post-cue 300-500ms, a larger pupillary dilation for exogenous attention was observed (endo (2-4) VS. exo (2-4): *t(57)*=-2.29, *p*=0.026, Cohen’s *d*=0.30).

The results remained largely unchanged when we dropped out the trials without perfect central fixations (see Methods, supplementary Fig. S1a and Fig. S2a-b). These findings suggested that exogenous attention would induce a higher level of alertness which was usually associated with larger pupil dilation.

### Experiment 2

To further identify the neural dynamics underlying the behavioral distinction between effects of the two types of retro-cues, experiment 2 was performed by another group of 20 participants with EEG. The design was identical to that in experiment 1 with only one exception for the layout of Gabor patches, where under the load-2 condition, the Gabor patches were bilaterally presented on both hemispheres of the screen to balance the visual inputs. One participant was excluded from behavioral analysis due to poor performance (see the above criterion) and another 4 participants were excluded from EEG analysis due to excessive artifacts in EEG signals. Again, behavioral results demonstrated a significant main effect of LOAD (*F*(1, 18) = 116.31, *p* =2.8e-9, η_p_^2^ = 0.87), a significant main effect of ATTENTION (*F*(2, 36) = 43.27, *p* =2.7e-10, η_p_^2^ = 0.71), and a significant LOAD*ATTENTION interaction (*F*(2, 36) = 7.10, *p* =0.003, η_p_^2^ = 0.28). Post hoc *t* tests further revealed an advantage of exogenous retro-cues only under the load-2 condition (Fig. 2b; 2_endo vs. 2_exo: *t(18)*=2.25, *p_adj_*=0.037, Cohen’s *d*=0.52; 4_endo vs. 4_exo: *t(18)*=0.06, *p_adj_*=0.952, Cohen’s *d*=0.01), replicating the findings in experiment 1. Furthermore, such benefit of the exogenous cue was not due to the speed-accuracy trade-off effect, as no reaction time (RT) difference was observed between the two types of retro-cues under any memory loads (see Supplementary Fig. S1b).

One lateralized event-related potential (ERP) component, an anterior directing attention negativity (ADAN, 300-500ms after cue onset) at frontal F3/4 electrodes, which was tightly related to attentional orientation (Eimer, Velzen, & Driver, 2002; Göddertz, Klatt, Mertes, & Schneider, 2018; Myers et al., 2015), showed a shorter latency for exogenous retro-cues than that for endogenous retro-cues under the load-2 condition (Fig. 3c-e, see supplementary Fig. S2e-f for un-subtracted bilateral waves; 2_endo vs. 2_exo: *t(14)*=2.20, *p*=0.045, Cohen’s *d*=0.57; 4_endo vs. 4_exo: *t(14)*=1.50, *p*=0.156, Cohen’s *d*=0.387). Interestingly, differences in ADAN amplitude were further positively correlated with differences between the two types of cues under the load-2 condition in Raw SD (*r*=0.71, *p*=0.003) (Fig. 3f). Although similar neural-behavioral correlations were found for the posterior contralateral negativity (PCN, supplementary Fig. S2), which is also associated with attentional orientation (Eimer et al., 2002; Göddertz et al., 2018; Heuer & Schubö, 2016; Luck & Hillyard, 1994; Myers et al., 2015), PCN failed to show the difference in latency between the two types of retro-cues.

In addition, lateralized early ERPs, such as P1pc and P1ac (100-160ms after the cue onset), were exclusively observed in exogenous retro-cue conditions under both memory loads (supplementary Fig. S2k and S2o), whereas the distractor positivity (Pd, another lateralized ERP, 580-680ms after cue onset), which has been linked to the inhibition of attentional shift (Schneider, Barth, Getzmann, & Wascher, 2017; Schneider, Mertes, & Wascher, 2016), was solely identified in endogenous retro-cue conditions (supplementary Fig. S2n).

### Experiment 3

Although the EEG recording has the advantage of high temporal resolution, it still lacks of spatial resolution, and is not well-performed in source analysis, which could be remedied by MEG recording. Therefore, experiment 3 was performed by another group of 27 participants in a magnetoencephalography (MEG) scanner. As in the perceptual domain, when the cue validity drops to 50%, effects induced by endogenous cues diminish while effects induced by exogenous cues remain (Briand, 1998; Kingstone, Smilek, Ristic, Friesen, & Eastwood, 2003; Posner, 1978). To exclude cue validity as a potential factor that could influence endogenous vs. exogenous cues differently, we added a condition of cue validity with 50% in experiment 3. Meanwhile, in order to draw similar number of trials as in experiments 1 and 2, we focused on the load-2 condition in which endogenous and exogenous cues induced behavioral difference in the above experiments. The cue validity was block-wise designed with 100% vs. 50%. The combination of retro-cue types (endogenous or exogenous cue) and cue validities (100% or 50%) was presented at separate blocks, randomly intermixed with no-cue trials (see Methods for details).

Two subjects were excluded due to poor performance. The behavioral results revealed a significant main effect of ATTENTION when cues were 100% valid (*F*(2, 46) = 12.24, *p* =5.5e-5, η_p_^2^ = 0.35). Post hoc *t* tests further showed that exogenous cues led to lower Raw SD compared to endogenous cues (endo vs. exo: *t(23)*=2.20, *p_adj_*=0.038, Cohen’s *d*=0.45), consistent with results in experiments 1 and 2. In contrast, non-significant effect of ATTENTION was observed when cues were 50% valid (Fig. 2c; *F*(2, 46) = 1.14, *p* =0.329, η_p_^2^ = 0.047), although valid cues resulted in better performance than invalid cues (supplementary Fig.S1c; valid_endo vs. invalid_endo: *t(23)*=-2.58, *p*=0.017, Cohen’s *d*=0.53; valid_exo vs. invalid_exo: *t(23)*=-3.46, *p*=0.002, Cohen’s *d*=0.71). These results were different from the traditional findings in the perceptual domain, in which exogenous cues always attracted attention even when they were uninformative (Briand, 1998; Carrasco, 2011; Emmanuel Guzman, Marcia Grabowecky, German Palafox, 2012; Kingstone et al., 2003; Posner, 1978), suggesting that the differences between the two types of retro-cues in our study was not due to automatic attentional processing. Again, the control analysis on RTs in experiment 3 excluded the possibility of speed-accuracy trade-off (supplementary Fig. S1d). Taken together, there was a consistent behavioral difference between endogenous and exogenous retro-cues with 100% validity under the load-2 condition.

Another two subjects were excluded for MEG analysis due to large artifacts in MEG signals (see Methods for details). To assess brain areas that might lead to behavioral changes, we estimated neural responses in the source space for each condition. Two clusters including the right intraparietal sulcus (IPS) and left DLPFC displayed significantly larger neural activation in both cued conditions *vs.* no-cue condition in the post-cue period of 0-500 ms (permutation test <0.05). During the post-cue period of 100-300ms, activations at the left DLPFC (lDLPFC) showed a shorter latency for the exogenous cue than that for the endogenous cue (Fig. 3h-i; endo vs. exo: *t(21)*=2.20, *p*=0.040, Cohen’s *d*=0.47). Furthermore, during the post-cue period of 300-500ms which overlapping the time window of ADAN, differences in activation between endogenous and exogenous cues at the lDLPFC were negatively correlated with their behavioral differences in Raw SD (*r*=-0.55, *p*=0.008) (Fig. 3j). These findings, together with the ADAN in experiment 2 (Fig. 3g), further suggested that although in the perception domain, exogenous cues were regarded as merely bottom-up manner (Buschman & Miller, 2007; Hahn et al., 2006; Theeuwes, 2010), they induced a faster and stronger lDLPFC activation, which accounted for better performance than that induced by endogenous retro-cues. In addition, we found that activations at right IPS (rIPS) during the post-cue 100-300ms was marginally faster for exogenous than endogenous cues (supplementary Fig. S3a-b; endo vs. exo: *t(21)*=2.06, *p*=0.052, Cohen’s *d*=0.44), and could predict both retro-cue benefits in Raw SD (supplementary Fig. S3c-d; endo-neu: *r*=-0.55, *p*=0.008; exo-neu: *r*=-0.60, *p*=0.003), suggesting shared neural mechanisms by the two types of retro-cues in the parietal cortex.

Indeed, the superior parietal lobe (SPL) *vs.* the inferior parietal lobe (IPL) have been regarded as critical regions in top-down *vs.* bottom-up attention-related processes (Ciaramelli, Grady, & Moscovitch, 2008; Corbetta, Kincade, & Shulman, 2002; Corbetta & Shulman, 2002). To understand how visual information was transferred across parietal and frontal regions and how such process was modulated by frontal activities relating to behavior, Granger causality analysis was performed to the source activity of SPL, IPL, DLPFC, and the lateral occipital cortex (LOC) (see Methods for details, supplementary Fig. S3e). Results revealed that both endogenous and exogenous cues induced a top-down control from DLPFC to LOC during 300-500ms after the cue onset (Fig. 4a-b). The time window coincided with that of the frontal-behavioral correlation in experiment 3 (Fig. 3j), the pupil size difference in experiment 1 (Fig. 3b) and ADAN component in experiment 2 (Fig. 3f). Interestingly, such process exclusively took place as early as 200ms after the onset of an exogenous cue, consistent with the time window of lDLPFC activations in experiment 3 (Fig. 3h-i) and P1ac in experiment 2 (supplementary Fig. S2o). These findings further suggested that exogenous retro-cues induced early activation in the frontal-posterior network, which was quite different from the merely bottom-up processes induced by exogenous cues in perception (Buschman & Miller, 2007; Chica et al., 2013; Connor, Egeth, & Yantis, 2004; Dugué, Merriam, Heeger, & Carrasco, 2018; Eimer et al., 2002; Hickey, Van Zoest, & Theeuwes, 2010; Hopfinger & West, 2006; Theeuwes, 2010). For the no-cue condition, there was no information transfer between the frontal and occipital regions (Fig. 4c).

**Figure 4.**
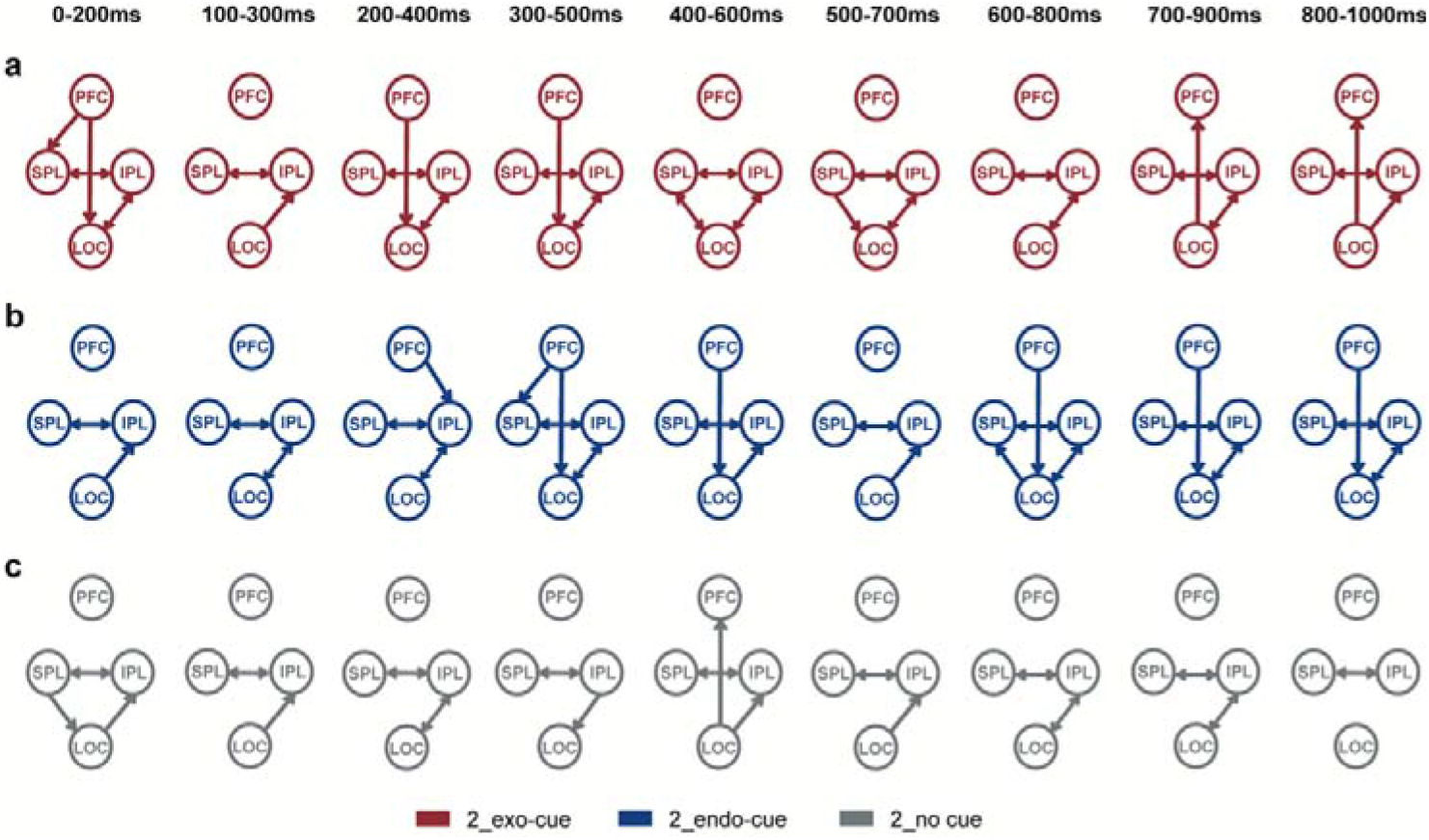
Granger causality analysis from experiment 3. **a-b**, Neural dynamics for exogenous (a) or endogenous (b) retro-cue trials with 100% validity, time-locked to the cue onset. **c,** Neural dynamics for no-cue trials in blocks with 100% validity, time-locked to the middle of delay period. Each arrow indicates the direction of information flow after Bonferroni correction (*p*<0.05).

### Experiment 4

Finally, we continued to identify the causal role of DLPFC and IPS in the retro-cue effects in experiment 4 with transcranial magnetic stimulation (TMS). A new group of 22 subjects were recruited. In each block, three types of cues were randomly mixed with equal number of trials. Stimulating sites were targeted at lDLPFC or rIPS, whose MNI coordinates were acquired as the maximal activation observed in experiment 3. Vertex served as a control stimulating site. To explore the time course of these brain regions in cognitive processes, a single pulse TMS was given at 100, 400, or 700ms after the cue onset (when no cue presented, we match the time points with the cued condition) in each trial. After considering the length of the experiment and maximum number of stimulating pulses in one session, we focused on the load-2 condition with 100% validity, in which all three experiments showed behavioral difference between endogenous and exogenous cues (see Methods for details).

Two subjects were excluded due to poor performance. When TMS was applied at vertex (Fig. 5a), there was a main effect of ATTENTION (*F*(2, 38) = 18.11, *p* =3.0e-6, η_p_^2^ = 0.49). Post hoc *t*-tests replicated the findings in the prior three experiments, indicating a superiority effect of exogenous retro-cues (endo vs. exo: *t(19)*=2.33; *p_adj_* =0.031; Cohen’s *d*=0.52). Interestingly, TMS over lDLPFC or rIPS both abolished the differences in Raw SD between the two types of retro-cues (lDLPFC: *t(19)*=-1.62, *p_adj_* =0.121, Cohen’s *d*=0.36; rIPS: *t(19)*=0.963, *p_adj_* =0.348, Cohen’s *d*=0.22).

**Figure 5.**
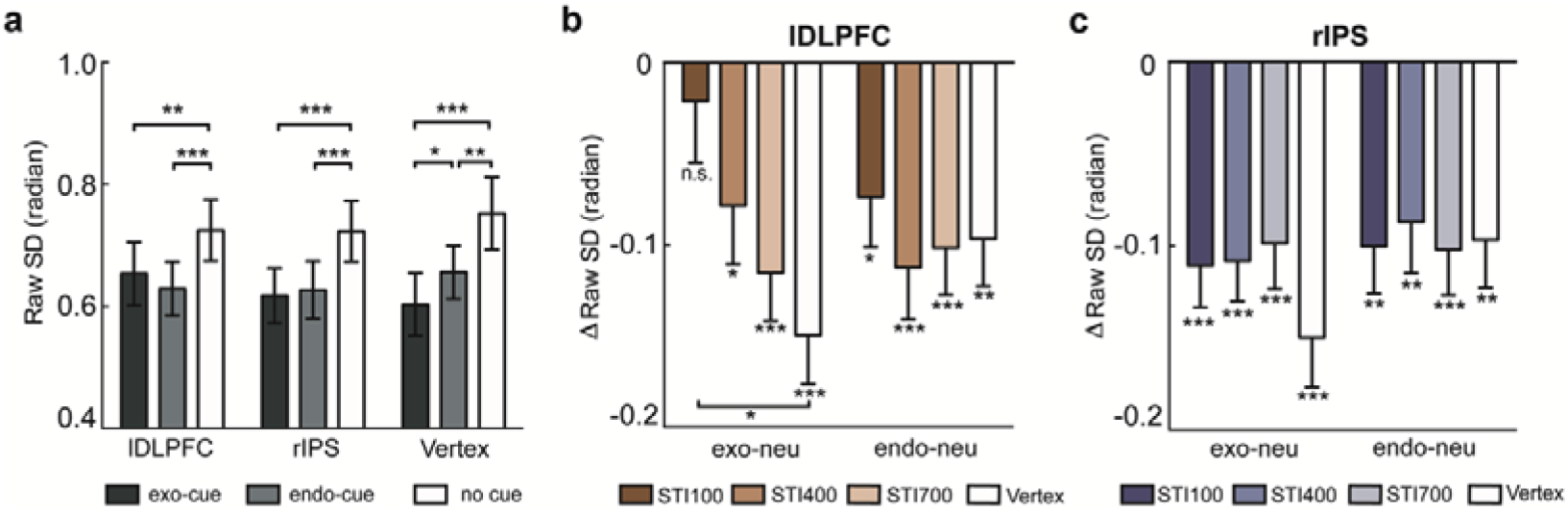
TMS results from experiment 4 (*n*=20). **a**, Merged data when TMS was set to LPFC, RIPS and vertex, respectively. **b-c,** Retro-cue effects when lDLPFC-targeted (b) or rIPS-targeted (c) TMS was delivered at 100ms, 400ms, and 700ms after the cue onset. ‘exo’ stands for exogenous cue, ‘endo’ stands for endogenous cue, and ‘neu’ stands for no cue condition. * *p*<0.05, ** *p*<0.01, *** *p*<0.001, all *p*-values are FDR corrected.

However, further looking at TMS results at different time points, they further suggested the timing of lDLPFC/rIPS function. When TMS was delivered at 100ms after the exogenous cue onset, the retro-cue benefit in Raw SD diminished (Fig. 5b; lDLPFC_100ms vs. vertex: *t(19)*=2.80, *p*=0.012, Cohen’s *d*=0.62). The control analysis on endogenous retro-cue effects at any stimulating time points did not replicate such results, neither did rIPS-targeted TMS at both types of retro-cues (Fig. 5b-c). These findings strongly proved that lDLPFC (but not rIPS) causally affected exogenous (but not endogenous) attentional processing within a time window of 100ms.

## DISCUSSION

Consistent with previous findings using retro-cues under the VWM load-4 condition (Kuo, Stokes, & Nobre, 2012; Murray et al., 2013; Pertzov et al., 2013; Pertzov, Manohar, & Husain, 2017; Shimi et al., 2014; Astle, Summerfield, Griffin, & Nobre, 2012; Gözenman, Tanoue, Metoyer, & Berryhill, 2014; Gunseli, van Moorselaar, Meeter, & Olivers, 2015; Lepsien & Nobre, 2006; Griffin & Nobre, 2003; Tanoue & Berryhill, 2012), we observed equivalent behavioral benefit for both exogenous and endogenous cues, confirming the important roles of spatial attention in modulating working memory representations. More importantly, we discovered a reliable difference in behavior between the two attentional effects under the low (but not high) VWM load through four experiments with 133 participants, which was not investigated in the previous studies (Kuo et al., 2012; Murray et al., 2013; Pertzov et al., 2013, 2017; Shimi et al., 2014; Astle, Summerfield, Griffin, & Nobre, 2012; Brady & Hampton, 2018; Gözenman, Tanoue, Metoyer, & Berryhill, 2014; Gunseli, van Moorselaar, Meeter, & Olivers, 2015; Lepsien & Nobre, 2006; Williams, Hong, Kang, Carlisle, & Woodman, 2013; Griffin & Nobre, 2003; Matsukura, Cosman, Roper, Vatterott, & Vecera, 2014; Tanoue & Berryhill, 2012). Combining multimodal neurophysiological recordings with eye-tracking, EEG, MEG and TMS, we further revealed convergent evidence for physiological and neural differences between the two attentional processes under the low VWM load.

Specifically, compared to endogenous cues, exogenous cues led to shorter latency of frontal ERP component (i.e. ADAN) indicating faster attentional deployment (experiment 2), and earlier DLPFC activity that were normally regarded as high-level processes of cognitive functions (experiment 3). Importantly, both amplitude of ADAN (experiment 2) and activity in lDLPFC (experiment 3) from 300ms to 500ms after the retro-cue onset explained the behavioral difference between the two types of cues.

How does such faster frontal activity in exogenous attention than endogenous attention come out and lead to better performance? Our data demonstrate one possibility. In experiment 2, there existed early contralateral ERP component (i.e. P1ac and P1pc) between 100-200ms after the exogenous cue, indicating fast automatic attentional processes, which however, did not exist in the endogenous condition. Such time window coincided with the time window in experiment 3 when the exogenous cues induced earlier DLPFC and IPS activity. Interestingly, the pupil dilation, which was tightly connected to the arousal systems (Alnæs et al., 2014; Granholm & Steinhauer, 2004; Nassar et al., 2012), appeared stronger for the exogenous cues than the endogenous cues at a similar latency of around 400ms (experiment 1). The overlapped time window of 300-500ms might indicate interactions between the alerting and orienting attention networks (Fan et al., 2005; Gazzaley & Nobre, 2012). To sum up, through the fast-automatic bottom-up processing, exogenous cues recruit early top-down control from DLPFC, and induce earlier frontal activities, which further interact with the arousal system that may amplify the processes. Such frontal activity induced by the exogenous cue is the real reason for the larger behavioral benefit than the endogenous cue.

However, it should be noted that both DLPFC and IPS demonstrate earlier activity in exogenous condition (experiment 3), and both the frontal ADAN and the parietal PCN could explain the behavioral difference between the two attentional types (experiment 2). Meanwhile, there are lots of evidence that prefrontal cortical areas receive bottom-up projection from parietal and occipital areas (Bressler, Tang, Sylvester, Shulman, & Corbetta, 2008; Connor et al., 2004; Theeuwes, 2010). Then which activity is more critical, the frontal or the parietal? In the current task-set, we suggested the more important role in the frontal top-down modulation, as TMS over DLPFC abolished the exogenous superiority but TMS over IPS did not (experiment 4). Nevertheless, it should be acknowledged that there might be other parietal areas that contribute to these processes as we only tested IPS in the present study.

Alternative explanations may exist for such top-down signal from DLPFC. For example, DLPFC has been suggested to maintain VWM representations in a plethora of studies (Barbey, Koenigs, & Grafman, 2013; Curtis & D’Esposito, 2003; S. Funahashi, Bruce, & Goldman-Rakic, 1989; Shintaro Funahashi, Chafee, & Goldman-Rakic, 1993; Shintaro Funahashi, Takeda, & Watanabe, 2004; Fuster, 1971). Recent studies reveal that the representations maintained in DLPFC neurons could be flexibly switched during the delay (Spaak, Watanabe, Funahashi, & Stokes, 2017; Stokes et al., 2013). Therefore, the top-down signal observed from the present study could also be representations stored in DLPFC and exogenous cues lead to faster shift of tuning of the representations (Spaak et al., 2017; Stokes et al., 2013). Indeed, dynamic coding in the prefrontal cortex has recently been regarded as the format of VWM storage (Meyers, 2018; Stokes, 2015), in contrast to the traditional view of static sustained activity (Goldman-Rakic, 1995). Our data put another possibility that attentional processes might also play a role in such dynamic shifting of representations.

Moreover, our data could unify attentional processes in the perceptual domain and those in the memory domain on the one side. In the current study, inspired by findings from the perceptual domain that the exogenous and endogenous attention effects tend to differentiate when sensory signal-to-noise ratio is high (Lu & Dosher, 1998, 2000; Lu, Lesmes, & Dosher, 2002), we employ a low load condition in which memory signal-to-noise ratio is high, as well as more sensitive measures with the memory recall task compared to the prior change detection task. Interestingly, we consistently find behavioral difference between exogenous and endogenous cues in four experiments. In such a way, the attentional effects in the perception domain and the memory domain can be incorporated by the same criteria of signal-to-noise ratio: endogenous and exogenous cues are equivalent when the signal-to-noise ratio is low; they tend to differentiate when the signal-to-noise ratio is high. However, future studies are needed to quantify the boundary of signal-to-noise ratio leading to the difference.

Meanwhile, at the neural level, contralateral ERP components indicating the attentional shift, such as ADAN and PCN, were identified for both endo- and exogenous retro-cues in the current study (experiment 2). Similar results were found in those EEG studies using spatial pre-cues (Eimer et al., 2002; Heuer & Schubö, 2016; Hickey, Lollo, & Mcdonald, 2009; Luck & Hillyard, 1994). Both endogenous and exogenous cues activated a large network involving parietal, frontal and visual cortices (experiment 3), overlapping the areas in fMRI studies using the pre-cues (Corbetta, Kincade, Ollinger, Mcavoy, & Gordon, 2000; Lepsien & Nobre, 2006; Mao, Zhou, Zhou, & Han, 2006; Rosen et al., 1992; Vandenberghe, Gitelman, Parrish, & Mesulam, 2001; Yantis et al., 2002; Dugué et al., 2018). These findings altogether indicated that spatial orienting in both the perceptual and VWM domain shared some common neural substrates.

On the other side, however, the attentional processes in perception or memory can be fairly different. Traditionally, exogenous attentional processes in perception were regarded as a merely bottom-up manner (Buschman & Miller, 2007; Chica et al., 2013; Connor et al., 2004; Dugué et al., 2018; Eimer et al., 2002; Hickey et al., 2010; Hopfinger & West, 2006; Theeuwes, 2010). However, in the present study, Granger causality analysis with source activity in MEG suggested a top-down signal from the prefrontal cortex as early as 100ms after the exogenous cue (experiment 3), and single pulse TMS over lDLPFC at the same 100ms post-cue time indeed abolished the exogenous retro-cue effects (experiment 4). These results indicated that 1) exogenous attention processes in memory were not only bottom-up; 2) the early top-down influence from the prefrontal cortex was critical for the superiority of exogenous attentional effects.

Besides the early top-down signal distinguishing exogenous attention processes in the memory domain from those in the perceptual domain, there still existed other differences between the attentional processes in the two domains, based on our results. For example, exogenous cues behaved differently in the present study from that in the perceptual domain when the cue validity was 50%. Previous studies suggested that in the perceptual domain exogenous cues always attracted attention in the condition with 50% validity while endogenous cues did not (Briand, 1998; Kingstone et al., 2003; Posner, 1978). However, in the current results, both of their retro-cue benefits had gone when the cues were 50% valid, although differences between valid and invalid trials still existed. Altogether, at least we could say the attentional processes in memory vs. perception are similar but not the same, and the endogenous vs. exogenous attention processes in memory are different from the traditional top-down *vs.* bottom-up dissection in perception.

As predicted from the perceptual domain, the presence of superiority of exogenous cues towards VWM representations may be generalized to long term memory (LTM). It is of great interest to further demonstrate the two different types of attention effects on LTM at different signal-to-noise ratio of memory trace. Such findings may also facilitate the educational implications that when the task is easier and memory signal is strong, exogenous cues might be more efficient than endogenous cues.

Although we combined different research modalities to form convergent evidence to strengthen, and further to explain the superiority of exogenous attention on VWM when the memory load was low, there still remained questions to be resolved. For example, after the exogenous cue presents, how is the cue processed in sensory cortices and then conveyed to the prefrontal cortex to trigger the top-down control signal? Or it may be conveyed directly from subcortical areas such as the superior colliculus or pulvinar, given such a fast time scale. To addressing these questions may require invasive neurophysiological studies, combining with computational models. We look forward to future studies using these tools to fully understand the interaction between attention and VWM, both lies in the center of human cognition.

## METHODS

### Subjects

Sixty-four (age=22.84 ± 2.75 years; 21 males), twenty (age=21.07 ± 2.69 years; 6 males) and twenty-two subjects (age=21.32 ± 2.50 years; 10 males) were recruited in experiments 1, 2 and 4, respectively. Twenty-seven subjects (age=21.43 ± 2.18 years; 14 males) participated in experiment 3. All had a normal or corrected-to-normal vision and had no history of psychiatric or neurological disorders. Written informed consent was provided by all participants prior to the experiment. Experimental protocols were approved by the local research ethics committee.

### Stimuli and experimental tasks

The procedure is shown in Fig.1. Each trial started with a central fixation dot for 500 ms, followed by a memory display for another 500 ms. The display consisted of two or four Gabor patches (radius 5°, contrast 100%, spatial frequency 2 cycles/degree) within an imaginary Cartesian coordinate system. Their orientations were chosen at random differed by at least 10° in each trial. The distance from the center of each Gabor patch to the axes was 6.6°. After a delay period lasting 2,000 ms, a probe with a randomly oriented Gabor patch was presented at the position that was early encoded. Participants were asked to recall the orientation of the mnemonic Gabor patch as precise as possible by rotating the probe using a mouse. In retro-cue trials, an endogenous retro-cue (a white arrow at the center, 1.8° by 1.28°) or an exogenous retro-cue (a dashed white circle at the probed location, radius 5°) was presented for 100ms at the middle of the delay period. Following the retro-cue, a second delay period was presented for another 900ms. In no cue trials, the white fixation dot remained on the screen during the whole delay period, without any changes to it. The inter-trial interval was 1,000ms. At the beginning of the experiment, participants were instructed to maintain fixation on the central dot with minimum eye blinks during the first 3000ms in each trial.

In experiments 1, 2 and 4, the validity of retro-cues was always 100% correct, meaning that the probe always appeared at the cued position. Subjects were encouraged to make full use of the retro-cue (i.e., only covertly attend to the cued Gabor patch while ignoring the other Gabor patch(es)) in the retro-cue trials. Before the experiments, participants were requested to perform 20 practical trials. In both experiment 1 and 2, within-subject factors with cue types (endogenous cue *vs.* exogenous cue *vs.* no-cue) and memory loads (load-2 *vs.* load-4) were randomly mixed within each block. The whole experiment was divided into 8 blocks, each of which contained 72 trials.

In experiment 4, memory load was restricted to the load-2 condition, three cue types were randomly mixed within each block, each of which contained 72 trials. There were 23 blocks in total.

In experiment 3, one of the two retro-cue types (endogenous or exogenous) and its corresponding cue validity (100% correct or 50% correct) were manipulated in different blocks. These four types of blocks alternated for twice, and thus there were 8 blocks in total. Each block contained 80 trials, and four-fifth of them were retro-cue trials, leaving the no-cue condition randomly mixed in the remaining trials. Prior to experiment, subjects performed a brief practice (10 trials per block type) outside of the MEG scanner, following the same order as in the experiment to become familiar with the task. Block sequences were counterbalanced between subjects.

Notably, Gabor patches in the memory display were presented at any two of four quadrants under the load-2 condition in experiment 1, while in following three experiments, the two Gabor patches were bilaterally presented at the screen to keep the balance of visual inputs at both sides.

### Behavioral analysis

For each condition, the recall error was calculated by subtracting the response angle from the probe angle. Then, the standard deviation (Raw SD) of recall errors was calculated. Both the recall error and Raw SD values were adjusted for circular data by means of the CircStat toolbox for MATLAB® (Berens, 2009). Repeated measures ANOVA on Raw SD were conducted to identify the main effect of cue type (ATTENTION, experiment 1-4), memory load (LOAD, experiment 1-2), cue validity (experiment 3), stimulating position/time point (experiment 4), and their interactions. Adjusted p-values (p_adj_) were given by FDR correction when applying multiple comparisons. Partial eta squared (η_p_^2^) and Cohen’s *d* were given as measures of effect size.

In experiment 1, six participants were excluded due to poor performance (lower end in the group 99% confidence interval, the same below) in at least one condition or missing data, resulting in 58 subjects for further analysis. In experiment 2, only one subject was excluded from analysis due to poor performance. In experiment 3, Data acquisition for one subject was aborted upon the request of the participant. Another two subjects were excluded from data analysis due to poor performance. Thus, we analyzed behavioral data from 24 participants in experiment 3. In experiment 4, two out of twenty-two subjects were excluded from analysis due to poor performance.

### Pupil data analysis

In experiment 1, we used an Eyelink 1000plus eye (SR Research, Canada) to monitor the trajectory of eye movements. Data from each participant’s dominant eye was used. The pupil diameter was corrected by the baseline using the mean value of 100ms prior to the memory display on a trial-by-trial basis for each participant. To reduce the difference in pupil changes evoked by the differential sensory format of the cues, we focused on exploring the interaction between memory loads and cue types. Therefore, we conducted a 2-by-2 measures ANOVA with LOAD (load-2 *vs.* load-4) and ATTENTION (endogenous cue *vs.* exogenous cue) as independent variables at each time point. In a control analysis to eliminate the contamination from eye movements, good trials were selected off-line, in which fixation was maintained within 1.5° visual angle and no blink was observed throughout the first 3000ms in each trial. Finally, data from 34 participants were qualified and remained for further analysis, with at least 50 acceptable trials remaining in each experimental condition.

### EEG recording and preprocessing

In experiment 2, we recorded the brain activities using electroencephalography (EEG). EEG signals were recorded continuously from 32 Ag/AgCl active electrodes (Easycap; Berlin, Germany) according to the international 10/20 System (Pivik et al., 1993). Vertical electrooculogram (VEOG) was measured by an additional electrode applied below the right eye. A BrainAmp DC-amplifier (BrainProducts; Gilching, Germany) sampled EEG and VEOG signals with a frequency of 1000Hz. A 250Hz low-pass filter was used and the impedance of electrodes was kept below 5kΩ during recording.

EEGLAB (Delorme & Makeig, 2004) was mainly used for data analysis. All channels were re-referenced offline to the averaged mastoids by means of the signal recorded from electrodes TP9 and TP10. EEG data were filtered with both 0.5Hz high-pass and 40Hz low-pass filters, and then divided into segments ranging from 700ms before to 3000ms after the onset of the memory display. The mean value of 200ms prior to the memory display was served as the baseline on trial-by-trial basis. Epochs containing artifacts, such as blinks or saccades, and excessive noise (±75 μV) at any electrode within −200 and +2500ms time window were excluded from further analyses. After this operation, EEG data from 15 subjects were qualified with at least 50 trials in each condition. To obtain contra- and ipsilateral activities evoked by retro-cues, electrodes from both hemispheres were exchanged in trials where subjects were cued to recall the right side of the memory display. Then, the mean amplitudes across trials at each electrode were calculated. As a result, the curves from the right hemisphere stood for contralateral activities, and that from the left hemisphere stood for ipsilateral activities.

### ERPs analysis

ERPs were referred to the onset of retro-cues. Time ranges for analyzing PCN and Pd at posterior electrodes P7 and P8 were accordingly set to 350 – 450 ms and 580 – 680 ms, ADAN at anterior electrodes F3 and F4 were tested in the time window of 300 – 500 ms, in line with previous studies (Schneider, Barth, Getzmann, & Wascher, 2017; Schneider, Mertes, & Wascher, 2016). Posterior P1pc and anterior P1ac were identified when amplitudes approached their peaks within time ranges for P1 (100-160ms in our study). For statistical analyses, we conducted 2-by-3-by-2 repeated measures ANOVAs with LOAD (load-2 *vs*. load-4), ATTENTION (endogenous cue *vs*. exogenous cue *vs*. no cue) and hemispheres (contralateral *vs*. ipsilateral to cued side) as within-subject factors. Lateralized effects were further ensured using paired *t*-tests (i.e. contralateral *vs*. ipsilateral to the cued side) for each condition. To investigate lateralized differences between two types of retro-cues, paired *t*-tests were calculated for each memory load with mean amplitudes or latencies of event-related lateralization (ERL) within the corresponding time window as dependent variables.

### MEG procedure and preprocessing

In experiment 3, anatomical MRI scans were always conducted after MEG recordings. A T1-weighted image was acquired with a Siemens 3-Tesla Prisma scanner (192 sagittal slices, 1mm thick, TR=2.53s, TE=2.98ms, flip angle=7°, FOV=22.4 cm *25.6cm) in 17 subjects or with a GE scanner (192 sagittal slices, 1mm thick, TR =6.64ms, TE=2.93ms, flip angle=12°, FOV=25.6cm *25.6cm) in another 10 subjects (out of 27). The cortex with 1 mm *1 mm *1 mm resolution was then extracted using the FreeSurfer toolbox, for estimating the source activity in the brain (Reuter, Schmansky, Rosas, & Fischl, 2012).

MEG data were obtained with a whole-head 306-channel Vector view system (Elekta-Neuromag, Helsinki, Finland), consisting of 102 magnetometers and 204 orthogonal planar gradiometers. The signal was recorded at a sampling rate of 1000Hz with an online bandpass filter from 0.1 to 250Hz. The head position was measured at the beginning of the experiment with six head position indicator coils. Anatomical landmarks (nasion, brow, left and right ear) and extra points (∼100) of the head shape were obtained using a 3D digitizer (Fastrak Polhemus, Colchester, VA, USA). External noise was removed with a signal space separation (SSS) method implemented with MAX filter software (Taulu, Kajola, & Simola, 2004). Head positions in the following runs were re-aligned to the position measured in the first run. During this step, MEG data from two subjects were further excluded due to a technical failure, thus we analyzed the remaining MEG data from 22 subjects. Further analyses were conducted with custom scripts based on Brainstorm toolbox (Tadel, Baillet, Mosher, Pantazis, & Leahy, 2011) for MATLAB®. Continuous data were notch filtered using 50, 100, and 150Hz filters, and then band-pass filtered between 1 and 100Hz. Muscular artifacts and eye blinks were evaluated with ICA, and were visually inspected and manually removed. Remaining data were then cut into 2600ms corresponding to 100ms before the memory display and 2500ms after the onset of memory display. Event-related field (ERF) analysis was conducted on 102 magnetometer channels.

### Source localization and selection of ROIs

MEG signals in the empty room, serving as covariance for noise modeling, were recorded prior to the formal experiment. We calculated the forward model using the method of overlapping spheres for each subject, and then applied the Minimum norm model (current density map) to estimate source activities for trials of each block. Next, neural activations at the source level for each condition were extracted and then averaged across trials before performing low-pass filter (below 32Hz) and z-score transformation. After this, absolute values for each condition from per subject were calculated and projected onto the MNI template. Two clusters including the right intraparietal sulcus (IPS) and left DLPFC displayed significantly larger neural activation in both cued conditions *vs.* no-cue condition in the post-cue period of 0-500 ms (permutation test α<0.05). To compare differences between two types of retro-cue conditions, paired *t*-tests were used with their latencies of brain responses within specific time windows (i.e., 100-300ms or 300-500ms after cue onset) as dependent variables. Pearson correlations between neural activations and behavior were applied to these brain regions, respectively.

### Granger causality analysis

It has been well documented that the superior parietal lobe (SPL) and inferior parietal lobe (IPL) are involved in voluntary and involuntary attentional processing, respectively (Corbetta & Shulman, 2002). To evaluate how the information from visual cortex interacted with these two regions, and also, how it received regulation from prefrontal cortex relating to behavior, the lateral occipital cortex (LOC) and dorsal lateral prefrontal cortex (DLPFC) were recruited for the analysis. To achieve this, we extracted time courses of these four brain regions, within a post-cue 1000ms time period, based on the Desikan-Killiany labeling system (Desikan et al., 2006). We then put them into the Multivariate Granger Causality (mvgc) toolbox (Barnett & Seth, 2014) to perform the Granger causality analysis, where a length of 200ms per time window stepped by 100ms was applied. Multiple comparisons were corrected using Bonferroni correction at a significant level of 0.05.

### TMS procedure

In experiment 4, anatomical T1-weighted magnetic resonance imaging (MRI) scans were always conducted preceding TMS sessions in a different day (interval >48h). They were acquired with a Siemens 3-Tesla Prisma scanner (192 sagittal slices, 1mm thick, TR=2.53s, TE=2.98ms, flip angle=7°, FOV=25.6 cm *25.6cm) at the ECNU MRI Research Center. These images were then imported into the BrainSight neuronavigation software (BrainSight 2.0, Rogue Research, Montreal, Canada) to allow for stereotaxic registration of the coil with the brain. TMS was delivered via Magstim Rapid2 stimulator and a 70-mm figure-of-eight coil (The Magstim Company, Whitland, UK). The TMS intensity for each subject was calculated as 90% of phosphene threshold (output= 55% ± 6.5%). The threshold was determined as subjects perceived phosphenes in 5 out of 10 TMS pulses given at the lateral occipital (LO) region, localized to be 1–1.5 cm caudal on the skull in a direct line towards the inion in accordance with various anatomical and functional maps (Van Essen et al., 2001).

During the experiment, TMS was applied to either the left dorsal lateral prefrontal cortex (lDLPFC, MNI coordinates: x=-29, y= 33.0, z=28.5) or the right intraparietal sulcus (rIPS, MNI coordinates: x=24.8, y=-64.2, z=41.7) in 20 blocks, whose MNI coordinates were acquired as the maximal activation observed in experiment 3. Their stimulation sequence was counterbalanced between subjects. Importantly, we also recruited vertex as a control area where behavioral performance was supposed to be unaffected by stimulation. TMS applied to vertex was always arranged at the 1st,12th (middle) and 23rd (last) blocks to match subject’ states in other blocks which may be influenced by the practice effect as well as the fatigue effect.

In each trial, a single pulse TMS was applied at one of three time points (100, 400 and 700ms after the (no) cue onset) in a pseudo-random order. We collapsed data with different stimulating time points when TMS was applied to vertex. Therefore, the lDLPFC- or rIPS-targeted stimulation condition (each cue type * each time point) consisted of 80 trials each, and the vertex-targeted stimulation condition contained 72 trials per cue type.

---

**Supplementary figure 1.**
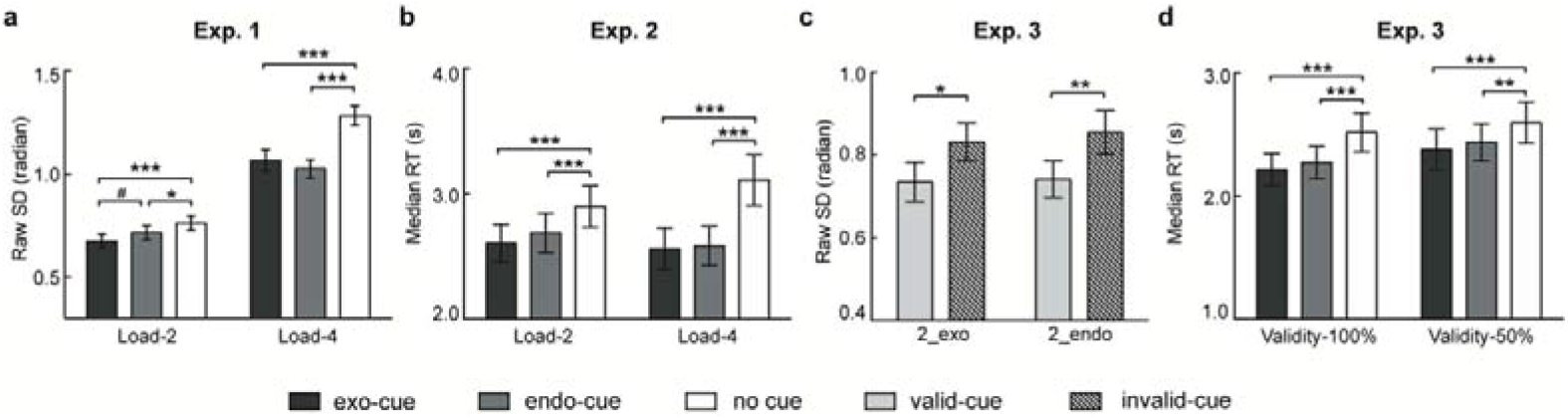
Behavioral results. **a**, Behavioral results from experiment 1 after controlling eye movements and blinks (*n*=34). **b,** Median reaction times (RTs) from experiment 2 (n=19). **c,** Behavioral results from experiment 3 for retro-cue trials with 50% cue validity (*n*=24). **d,** Median RTs from experiment 3. # *p*<0.1, * *p*<0.05, ** *p*<0.01, *** *p*<0.001, all *p*-values are FDR corrected.

**Supplementary figure 2.**
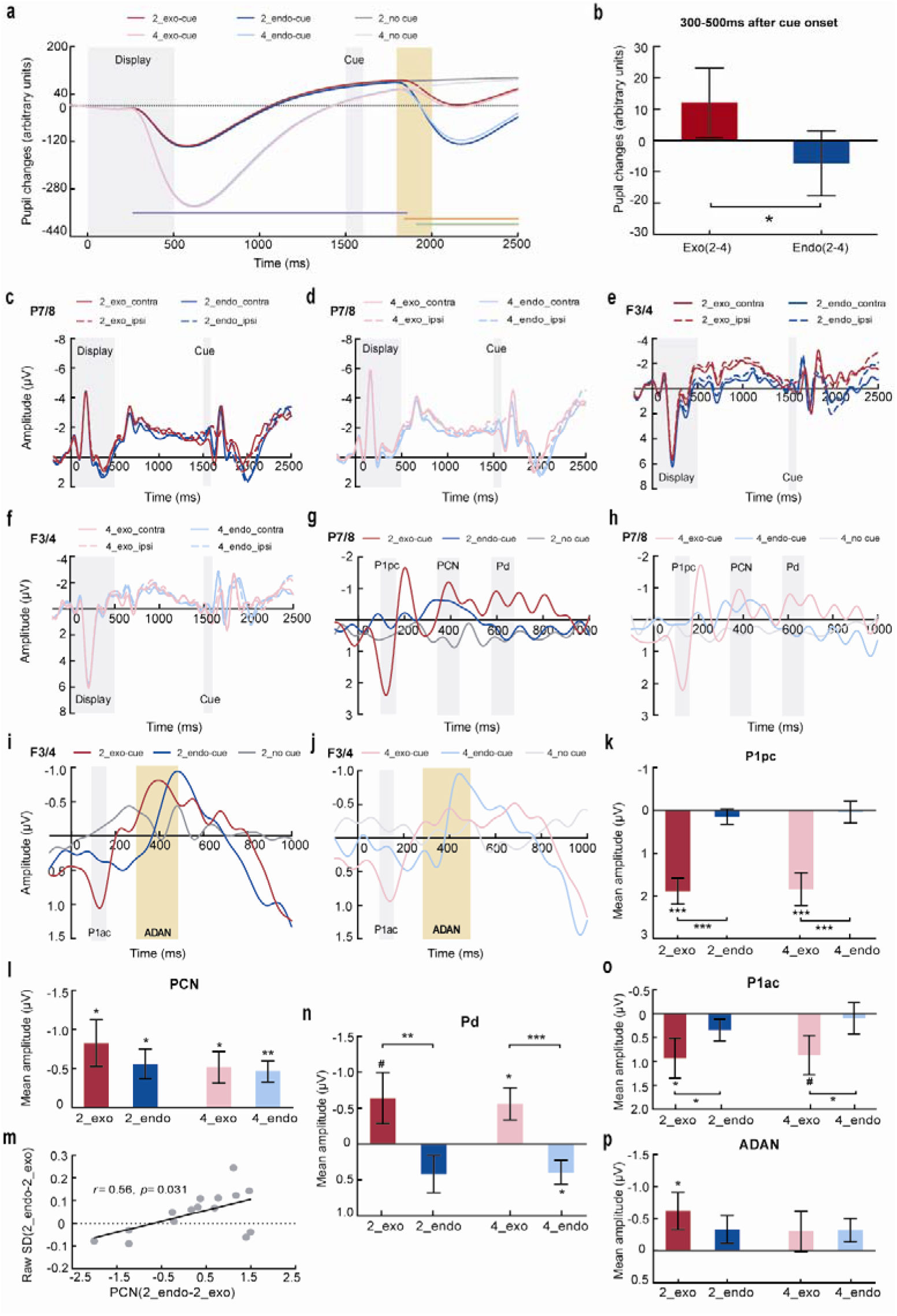
Psychophysiological results from experiment 1 and 2. **a**, Pupil changes using qualified trials from 34 subjects as a function of time referred to the memory display. Purple dots indicate a main effect of LOAD, orange dots indicate a main effect of ATTENTION, and green dots indicate a 2-by-2 interaction. **b,** Load-by-cue interactions within the time window of 300-500 ms after the cue onset. **c-f,** The original curves referred to memory display at P7/8 electrodes (c-d) and F3/4 electrodes (e-f). **g-j,** The contra-minus ipsilateral curves time-locked to the cue onset at P7/8 electrodes (g-h) and F3/4 electrodes (i-j). **k,** Mean amplitudes of P1ac. **l,** Mean amplitudes of PCN. **m,** Correlations between endo-minus exogenous PCN differences at the load-2 condition and their behavioral differences at the load. **n-p,** Mean amplitudes of Pd (n), P1ac (o), ADAN (p). # *p*<0.1, * *p*<0.05, ** *p*<0.01, *** *p*<0.001.

**Supplementary figure 3.**
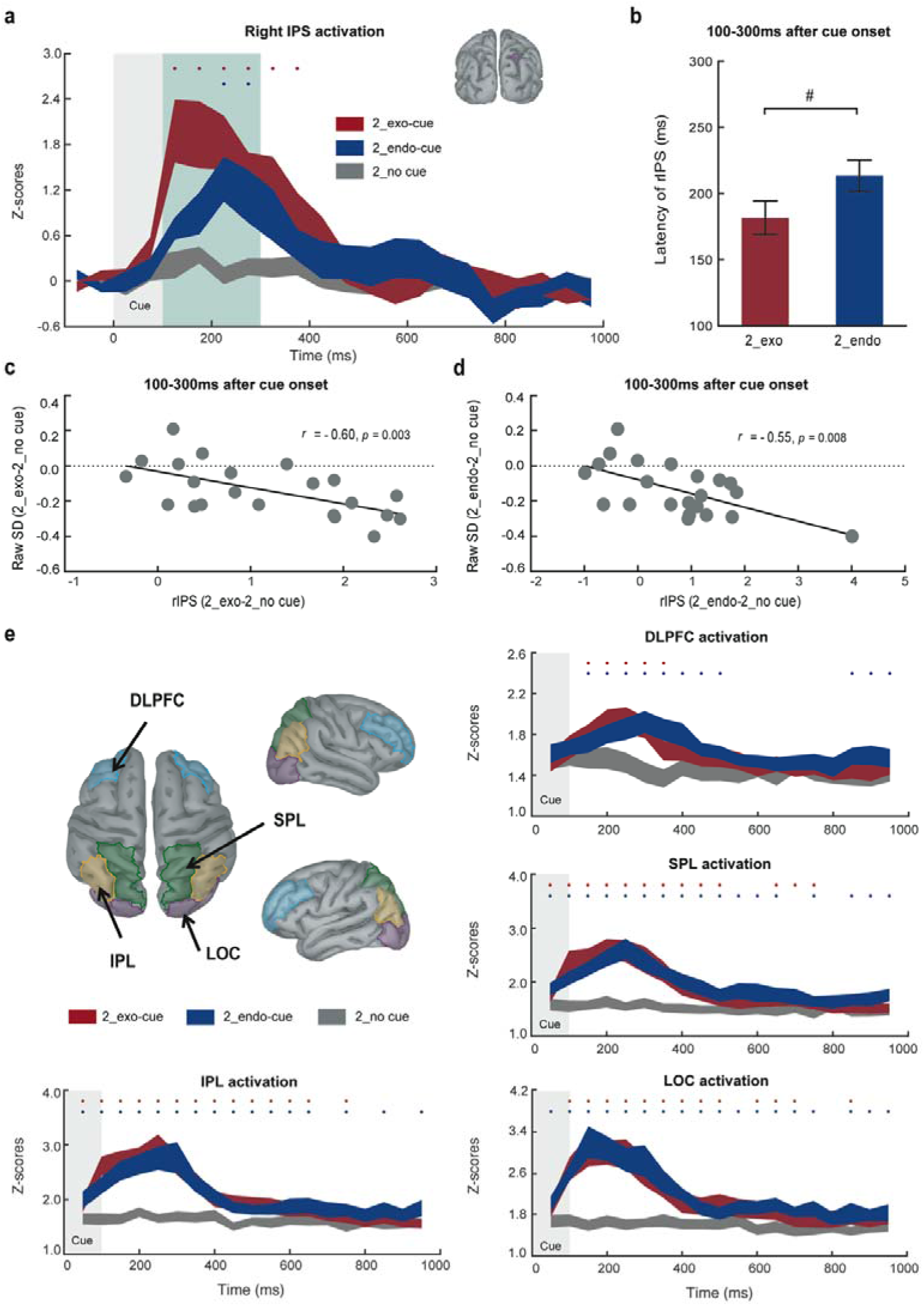
Psychophysiological results from experiment 3. **a-b**, Grand event-related field time-locked to memory display in blocks with 100% validity. Red (blue) dots indicate higher brain responses to exogenous (endogenous) retro-cues than to no cue condition (one-tailed *t*-tests, *p*<0.05). b, The latency of rIPS activation within the range of 100-300 ms after cue onset for retro-cue conditions. (# *p*<0.1). **c-d,** Correlations in blocks with 100% validity between the activation differences of exogenous (c) or endogenous (d) retro-cue relative to no cue conditions at rIPS and their behavioral differences within the time window of 100-300 ms after the cue onset (two-tailed *t*-tests, *p*<0.05). **e,** ROIs on a template and z-transferred grand responses within these ROIs in blocks with 100% validity time-locked to the cue onset. Red (blue) dots indicate higher responses to an exogenous (endogenous) retro-cue than to no cue (one-tailed *t*-tests, *p*<0.05).

